# Identification of essential oils with strong activity against stationary phase uropathogenic *Escherichia coli*

**DOI:** 10.1101/702951

**Authors:** Shuzhen Xiao, Peng Cui, Wanliang Shi, Ying Zhang

## Abstract

*Escherichia coli* is the most dominant pathogen causing urinary tract infections (UTIs), but the current most frequently prescribed antibiotics do not always effectively cure the infection due to quiescent persister bacteria. While it has been reported that some essential oils have antimicrobial activity against growing *E. coli*, the activity of essential oils against the non-growing stationary phase *E. coli* which is enriched in persisters has not been investigated. We evaluated the activity of 140 essential oils against stationary phase uropathogenic *E. coli* UTI89 and identified 39, 8 and 3 essential oils at 0.5%, 0.25% and 0.125% concentrations to have high activity against stationary phase *E. coli*. Among the top eight essential oils, Oregano showed higher activity than the known persister drug tosufloxacin. The other top seven hits included Allspice, Bandit "Thieves", Cinnamon bark, Syzygium aromaticum, Health shield, Cinnamon leaf and Clove bud. In Oregano essential oil drug combination studies with common UTI antibiotics, Oregano plus quinolone drugs (tosufloxacin, levofloxacin, ciprofloxacin) completely eradicated all stationary phase *E. coli* cells, partially enhanced the activity of nitrofurantoin, but had no apparent enhancement for fosfomycin, meropenem and cefdinir. Our findings may facilitate development of more effective treatments for persistent UTIs.

## 1. Introduction

Urinary tract infections (UTIs) are ranked as leading causes of healthcare-associated infections, which also account for significant morbidity and high economic costs [1, 2]. Among the pathogens causing UTIs, *Escherichia coli* is the most important one causing more than 75% of community-acquired UTIs and 30-50% of nosocomially-acquired UTIs [3]. Although there are some regimens recommended for treatment and management of UTIs [4], recurrent UTIs are reported in more than 40% of women who once had an UTI episode and thus pose a major clinical challenge [1]. While the cause for recurrent UTIs is debated, some studies support the theory that some persister bacteria are metabolically quiescent and tolerant to the current antibiotics [5, 6]. For instance, uropathogenic *E. coli* has been shown to form intracellular bacterial community (IBC) in bladder epithelial cells that is refractory to antibiotic treatment [7]. Despite the current antibiotic treatment, the quiescent persister bacteria are not killed effectively and can cause relapse upon withdrawal of antibiotics.

It is worth noting that the current antibiotics used in the treatment of UTIs such as nitrofurantoin, fosfomycin, sulfa drugs and quinolones are mainly active against the growing *E. coli* but have poor activity against the non-growing persisters [8]. In our previous study, we have screened an FDA-approved drug library and found some active hits such as tosufloxacin and colistin to be highly active against stationary phase culture of uropathogenic *E. coli* UTI89 strain [8]. The importance of including drugs that target persister bacteria has been demonstrated with persister drug colistin in a persistent UTI mouse model [9]. However, the choice of persister drugs that may be useful is quite limited from the FDA drug library screen [8]. In an attempt to identify additional drug candidates that could be useful for treating persistent UTIs, in this study, we evaluated a panel of 140 essential oils for their activity against stationary phase *E. coli* UTI strain UTI89. While some essential oils were found to be active against growing *E. coli* in vitro in previous studies [10–13], the number of essential oils evaluated is small (one to three essential oils) [10–13], and their activity against stationary phase cultures enriched in dormant persister bacteria have not been studied. The aim of this study is to evaluate the activity of a panel of essential oils against the stationary phase uropathogenic *E. coli* and identify active hits that may be useful for more effective treatment of persistent UTIs.

## 2. Results

### 2.1. Identification of active essential oils against stationary phase E. coli

Consistent with our previous studies [8], tosufloxacin included as a persistger drug control was shown to have high activity against stationary phase *E. coli*, while other clinical drugs including levofloxacin, ciprofloxacin, nitrofurantoin, sulfamethoxazole, trimethoprim, meropenem, cefdinir and fosfomycin were not able to completely kill stationary phase *E. coli* at 50 µM after three-day drug exposure [8]. Interestingly, after one-day exposure, 31 (Health Shield, Bandit "Thieves", Cinnamon bark, Syzygium aromaticum, Oregano, Clove bud, Allspice, Tea tree, Geranium bourbon, Lavender, *Cymbopogon flexuosus*, Marjoram, Peppermint, *Cymbopogon citratus*, Alive, Cumin, Deep muscle, Ho wood, Head ease, Litsea cubeba, Palmarosa, Thyme white, Caraway, Coriander oil, Dillweed, Neroli, Palo santo, Pennyroyal oil, Rosewood oil, Tarragon, Geranium Egypt), 5 (Health Shield, Bandit "Thieves", Cinnamon bark, Syzygium aromaticum, Oregano) and 3 (Oregano, Allspice, Cinnamon bark) essential oils were found to have high activity against stationary phase *E. coli* at 0.5%, 0.25% and 0.125% concentrations, respectively, with no growth detected on LB plates. When the drug exposure was extended to three days, an additional 8 (Cinnamon leaf, Spearmint, Cajeput, Citronella, Birch, Cornmint, Lemon eucalyptus, Bay oil) and 3 (Clove bud, Allspice, Cinnamon leaf) essential oils were found to be active at 0.5% and 0.25% concentrations (Table 1). The top 8 essential oils (Health shield, Cinnamon leaf, Clove bud, Bandit "Thieves", Cinnamon bark, Syzygium aromaticum, Oregano, and Allspice) which showed high activity at 0.25% concentration were used in the subsequent testing to confirm their activity in inhibiting growing *E. coli* in MIC test and in CFU drug exposure assay for their activity against non-growing stationary phase *E. coli*.

**Table 1.**
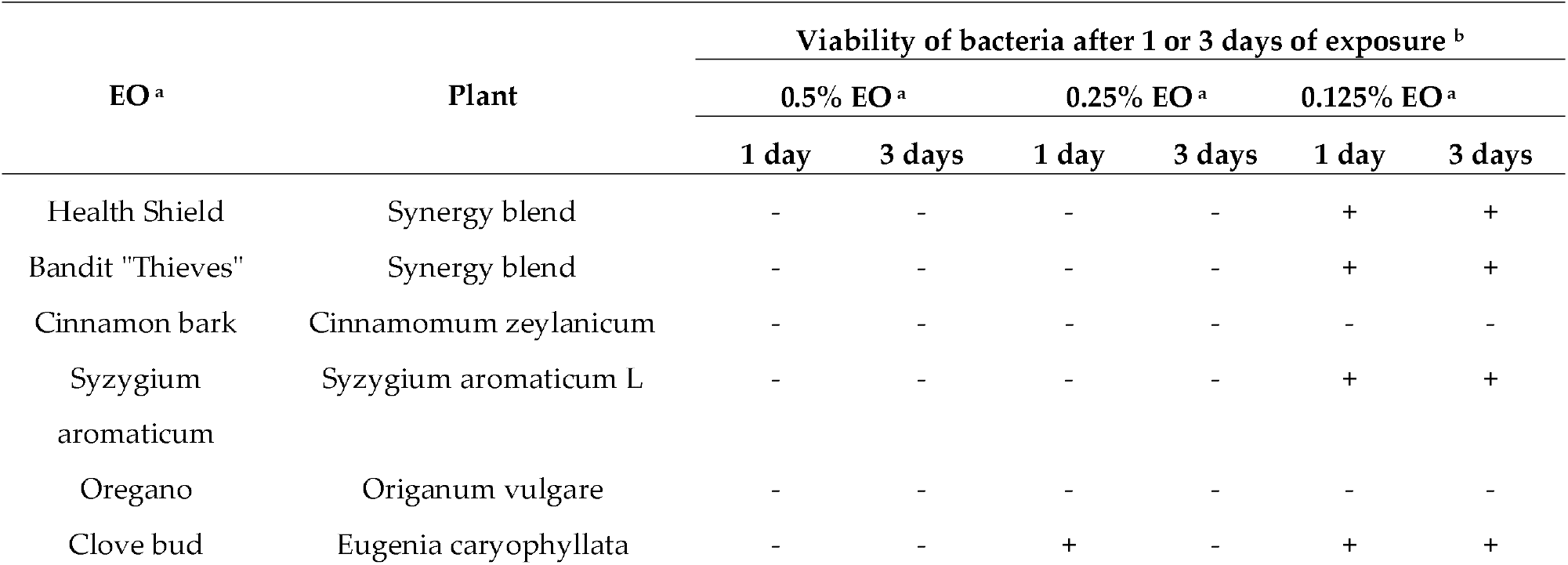
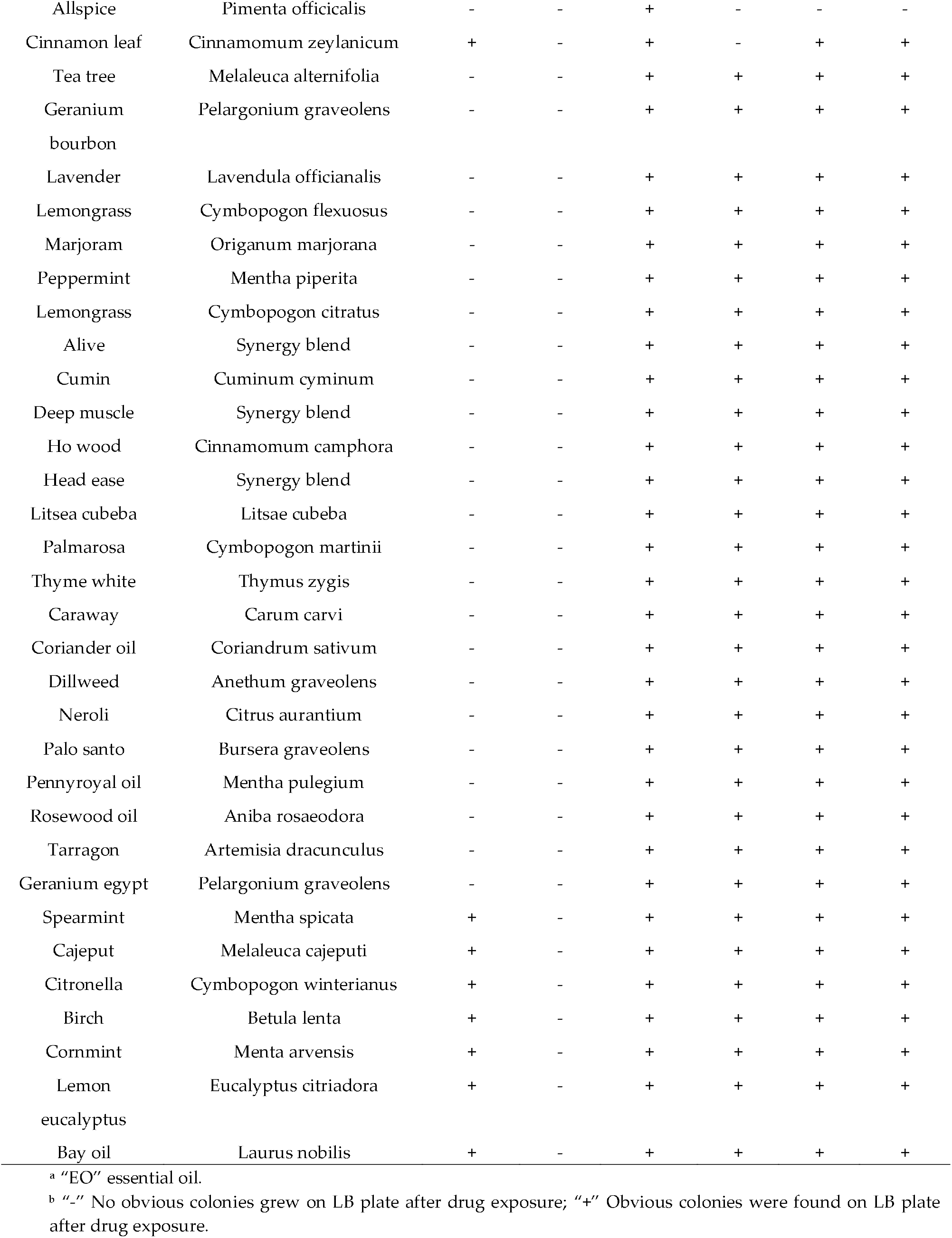
Effect of essential oils on stationary phase uropathogenic *E. coli* UTI89

### 2.2. MIC determination of the top active essential oils

We carried out antibiotic susceptibility testing to determine the activity of the top 8 active essential oils against growing *E. coli*. As shown in Table 2, Oregano was the most effective agent in inhibiting *E. coli*, with the lowest MIC (0.015%) in our study. The growth of *E. coli* was efficiently suppressed by Cinnamon bark at the concentration of 0.03%. While the other six essential oils (Health shield, Cinnamon leaf, Clove bud, Bandit "Thieves", Syzygium aromaticum and Allspice) were active with the same MIC value of 0.125%. Clinical drug fosfomycin included as a control inhibited the growth of *E. coli* growing cells with MIC of 16 µg/mL.

**Table 2.** Activity of top 8 essential oils that are active against stationary phase uropathogenic *E. coli* UTI89 in terms of their activity against growing bacteria (MIC) and non-growing bacteria in drug exposure

| Drug (50 µM)/essential oil (0.25%) | Plant | MIC (µg/mL /%) | Viability of bacteria after 1 or 3 days of drug exposure<sup>b</sup> |  |
| --- | --- | --- | --- | --- |
|  |  |  | 1 day | 3 days |
| Fosfomycin | - | 16 | - | - |
| Health shield | Cassia, clove, eucalyptus, lemon and rosemary | 0.125 | - | - |
| Cinnamon leaf | Cinnamomum zeylanicum | 0.125 | + | - |
| Clove bud | *Eugenia caryophyllata* | 0.125 | + | - |
| Bandit "Thieves" | cloves, cinnamon, lemon, rosemary and eucalyptus | 0.125 | - | - |
| Cinnamon bark | Cinnamomum zeylanicum | 0.03 | - | - |
| *Syzygium aromaticum* | *Syzygium aromaticum* L | 0.125 | - | - |
| Oregano | *Origanum vulgare* | 0.015 | - | - |
| Allspice | *Pimenta officinalis* | 0.125 | + | - |

<sup>b</sup> "-," No obvious colonies grew on LB plate after drug exposure; "+" Obvious colonies were found on LB plate after drug exposure.

### 2.3. Comparison of active essential oils in their abilities to kill stationary phase *E. coli* UTI89

According to our previous study [8], tosufloxacin had good anti-persister activity against uropathogenic *E. coli*. While levofloxacin, nitrofurantoin, sulfamethoxazole, trimethoprim and fosfomycin are widely used for treating urinary tract infections, they have poor or no obvious activity against uropathogenic persisters [8, 14]. In this study, we tested the activity of tosufloxacin and other clinical drugs against stationary phase *E. coli* at 20 µM. As previously described [8], tosufloxacin could kill all stationary phase *E. coli* cells after three-day drug exposure, with no visible colonies on LB plate. Levofloxacin and ciprofloxacin had weak activity with 10^5^~10^6^ CFU/mL bacterial cells still remaining after five-day exposure. In contrast, other clinical drugs including nitrofurantoin, sulfamethoxazole, trimethoprim, meropenem, cefdinir and fosfomycin did not show obvious activity against stationary phase *E. coli* even when the drug exposure was extended to five days (Figure 1). Meanwhile, 8 essential oils were found to have stronger activity than tosufloxacin (20 µM) at 1% concentration, with 100% clearance after just one-day exposure. At 0.5% concentration, five essential oils (Oregano, Cinnamon bark, Syzygium aromaticum, Allspice and Bandit "Thieves") could eradicate all stationary phase cells after one-day exposure. The other three essential oils (Health shield, Cinnamon leaf, Clove bud) could not clear all the cells after five-day exposure. At a lower concentration of 0.25%, we noticed that Oregano still exhibited strong activity against stationary phase *E. coli*, and no CFU could be found after one-day exposure (Figure 2). Meanwhile, Allspice could eradicate stationary phase *E. coli* cells after three-day exposure. On the other hand, Bandit "Thieves", Cinnamon bark and Syzygium aromaticum could eradicate stationary phase *E. coli* after five-day exposure. Health shield, Cinnamon leaf and Clove bud still could not wipe out the stationary phase *E. coli* culture after five-day exposure.

**Figure 1.**
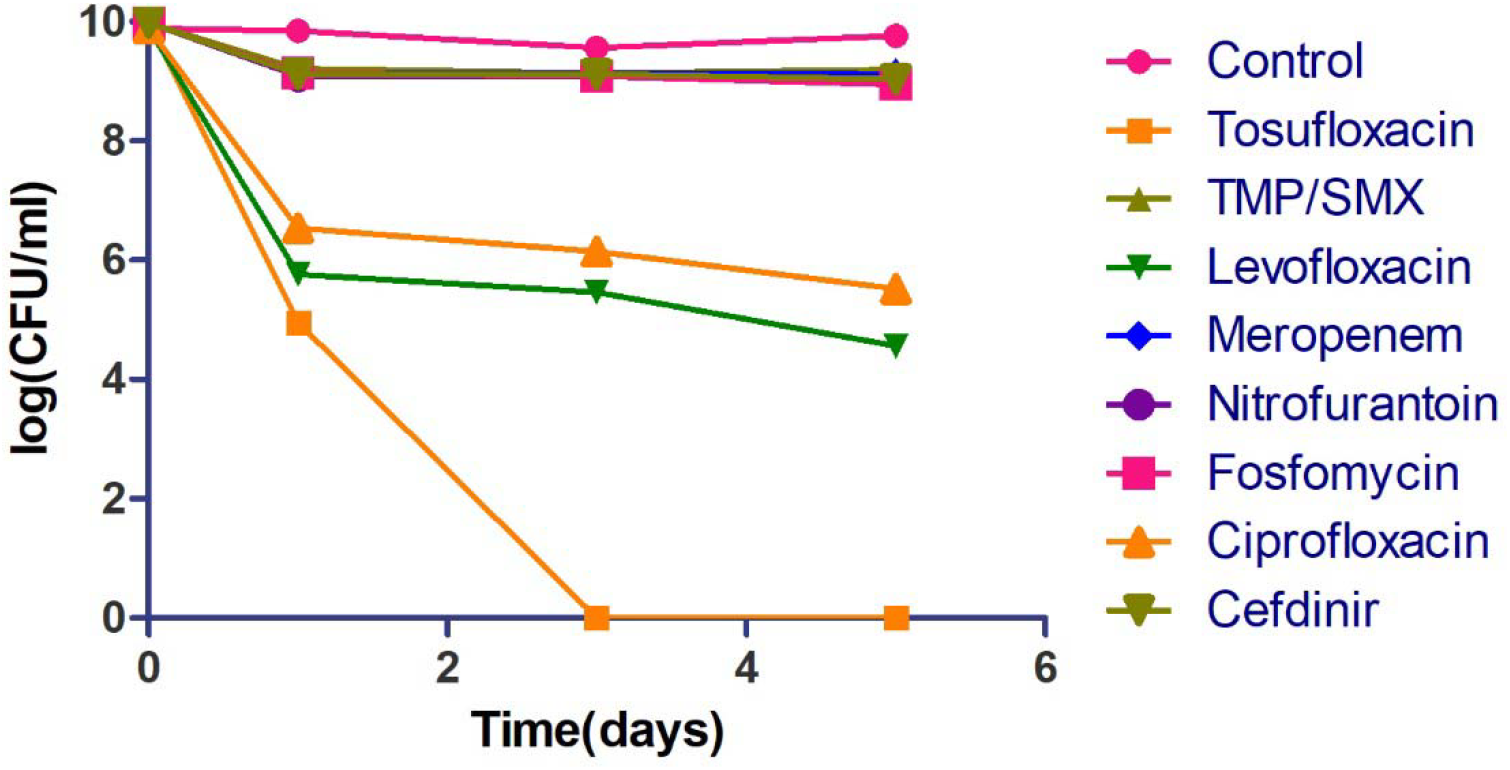
Activity of tosufloxacin and commonly used antibiotics against stationary phase *E. coli*. Tosufloxacin had good anti-persister activity against uropathogenic *E. coli*. Antibiotics commonly used to treat UTIs had poor activity against the stationary phase *E. coli*. The final concentration of antibiotics including tosufloxacin, levofloxacin, ciprofloxacin, nitrofurantoin, sulfamethoxazole-trimethoprim, meropenem, cefdinir and fosfomycin, was all 20 µM. Sulfamethoxazole-trimethoprim is the combination of trimethoprim and sulfamethoxazole in a ratio of 5:1.

**Figure 2.**
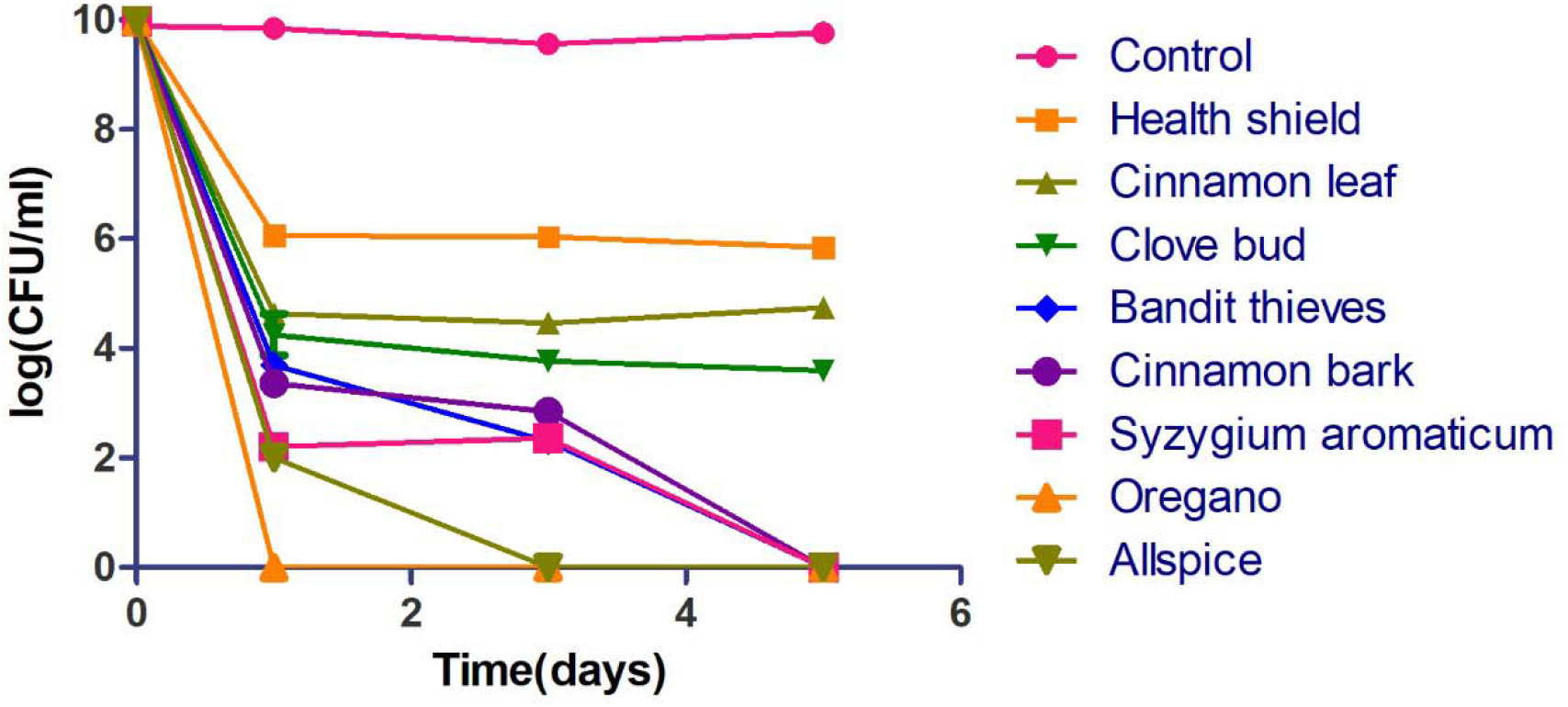
Activity of active essential oil candidates (0.25%) against stationary phase *E. coli*. Oregano could eradicate all stationary phase cells after one-day oil exposure. Allspice could kill all cells after three-day exposure. Bandit "Thieves", Cinnamon bark and Syzygium aromaticum could eradicate stationary phase *E. coli* after five-day exposure. Health Shield, Cinnamon leaf and Clove bud could not wipe out the stationary phase *E. coli* culture even after five-day exposure.

### 2.4. Development of essential oil drug combinations to eradicate stationary phase *E. coli* in vitro

According to our previous research [8] and this study, clinical drugs used to treat UTIs had limited activity to kill *E. coli* persisters. To more effectively eradicate the stationary phase *E. coli*, essential oil drug combinations were studied using clinical drugs in combination with Oregano (0.02%). The results showed that some drug combinations were indeed much more effective than single drugs (Figure 3). Among them, quinolone drug essential oil combinations including tosufloxacin + Oregano, levofloxacin + Oregano and ciprofloxacin + Oregano could completely eradicate all the stationary phase *E. coli* after seven-day exposure. Meanwhile, nitrofurantoin + Oregano combination killed the majority of stationary phase cells with 10^3^ CFU/mL cells remaining, showing much better activity than single drug nitrofurantoin (10^9^ CFU/mL cells remaining) and somewhat better activity than single Oregano (10^5^ CFU/mL remaining). In contrast, other drug combinations such as sulfamethoxazole + Oregano, trimethoprim + Oregano, meropenem + Oregano, cefdinir + Oregano and fosfomycin + Oregano had less activity against stationary phase cells, with 10^5^ CFU/mL bacterial cells remaining even after seven-day exposure, suggesting these combinations were not significantly better than Oregano alone (10^5^ CFU/mL remaining).

**Figure 3.**
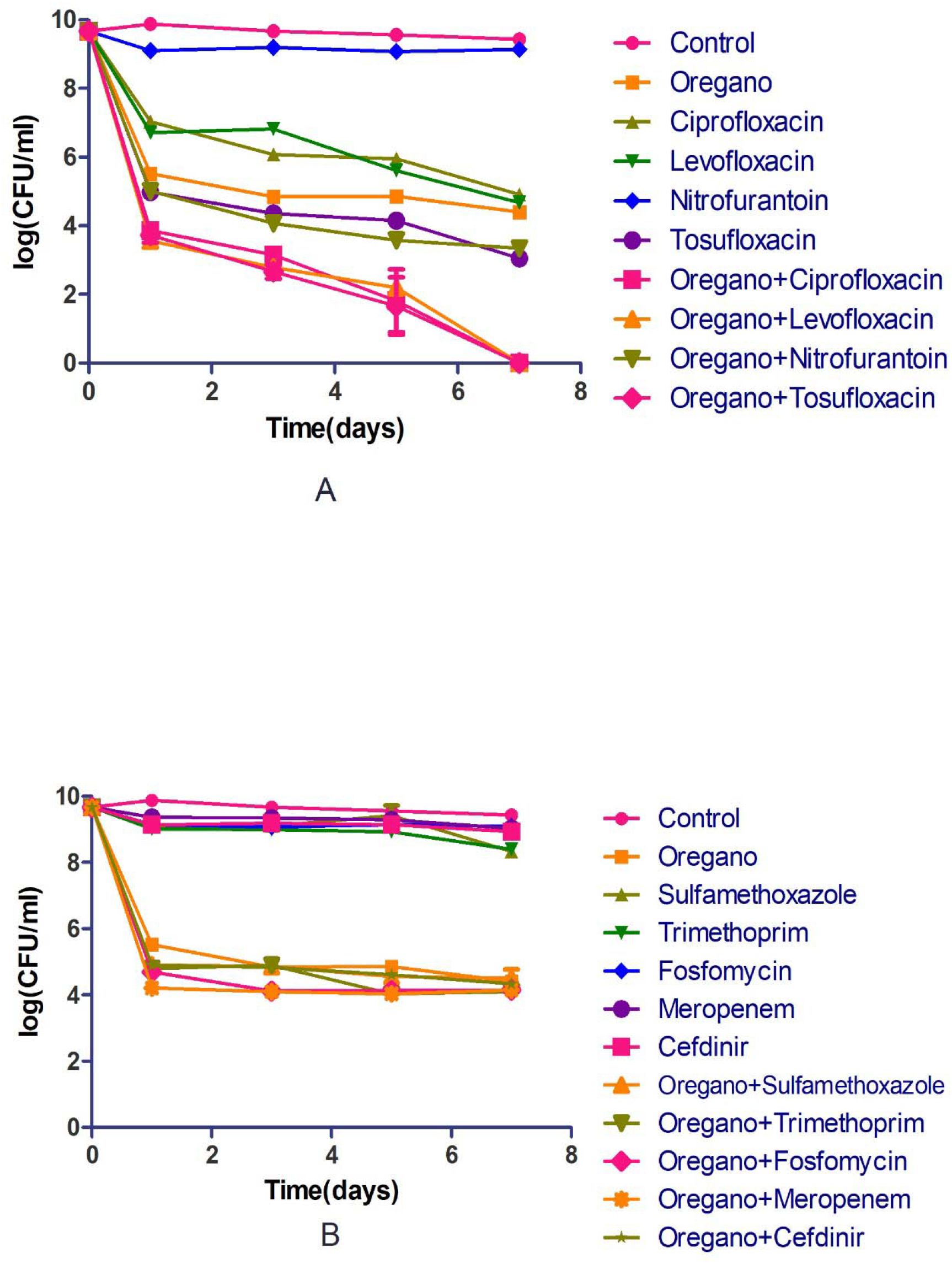
Comparison of the activity of Oregano with different antibiotic combinations against stationary phase *E. coli* UTI89 over time. Effects of ciprofloxacin, levofloxacin, tosufloxacin, nitrofurantoin and their combinations with Oregano are presented in (A). Effects of Sulfamethoxazole, trimethoprim, fosfomycin, meropenem, cefdinir and their combinations with Oregano are presented in (B). The final concentration of antibiotics is 5 µg/mL and the concentration of Oregano is 0.02%.

## 3. Discussion

The hypothesis that recurrent UTIs is primarily caused by persister clones is now widely accepted [5, 6]. While commonly used antibiotics for UTIs exhibited effective inhibitory effect on multiplying and free-living log phase *E. coli*, they demonstrated very limited activity against aggregated *E. coli* at stationary phase [8, 15, 16]. Our previous study with persister drug colistin in combination with gentamicin or fluoroquinolone drug indicated the importance of including persister drugs for more effective treatment of persistent UTIs in the mouse model [9]. In line with this thinking, in this study, we screened a panel of essential oils as suitable alternatives for more effective elimination of persister bacteria for improved treatment of recurrent UTIs.

Essential oils are concentrated volatile liquids that are extracted from plants, most of which are used as spices, culinary herbs and muscle relaxant [17, 18]. It has been shown that some essential oils have antimicrobial activity against log phase *E. coli* [11–13] and outstanding activity against stationary phase B. burgdorferi as shown in our previous studies [18, 19]. In this study, we identified 8 essential oils (at 1% concentration) that are more active than persister drug tosufloxacin (20 µM), a quinolone drug control that could eradicate stationary phase *E. coli* (Figure 1). Among them, five essential oils (Oregano, Cinnamon bark, Syzygium aromaticum, Allspice and Bandit "Thieves") exhibited remarkable activity against stationary phase *E. coli* at 0.25% concentration (Figure 2).

Oregano was used for gastrointestinal, diabetes, respiratory conditions, as antibacterial, anti-inflammatory in traditional medicine [20]. Its antimicrobial activity against *E. coli*, *Listeria innocua, Pseudomonas sp., and Salmomella sp.*, was confirmed in previous studies [21–23]. In this study, Oregano demonstrated high activity against not only log phase growing *E. coli* with a low MIC of 0.015% but also stationary phase non-growing bacteria with complete clearance at 0.25% concentration. Remarkably, in the study, we conducted the first in vitro drug combination study using a low concentration of Oregano with the currently recommended UTIs antibiotics including levofloxacin, ciprofloxacin, nitrofurantoin, sulfamethoxazole, trimethoprim, meropenem, cefdinir and fosfomycin. We found it was more effective to kill *E. coli* persisters by drug combination than single antibiotic, especially when Oregano was combined with levofloxacin and ciprofloxacin. When combined with tosufloxacin, another persister active drug, the combination also showed outstanding activity with 100% clearance after seven-day exposure. These finding may have implications for improve treatment of UTIs. Further studies are needed to validate such combination approaches are useful in animal models. Carvacrol has been identified as the main active component of oregano oil and is known to render the *E. coli* cell membrane permeable by causing loss of cell contents [10] and induce membrane damage in *P. aeruginosa* and *S. aureus* [24]. Further work should be carried out to confirm the active ingredient against persister *E. coli* as carvacrol and its mechanism of action.

Additionally, we found Cinnamon bark, Syzygium aromaticum, Allspice and Bandit "Thieves" showed excellent activity against stationary phase *E. coli*. Cinnamon bark has a long history in western medicine to soothe aching joint and numb pain and its bioactivity against bacteria, fungus, inflammation, cancer and diabetes has been described [25]. Here, we showed that Cinnamon bark oil had a low MIC (0.03%) and remarkable anti-persister activity. Syzygium aromaticum is one of the two plants (the other is Syzygium polyanthum) in Syzygium species that were reported to show antibacterial activity [26]. While its inhibitory effect on the growth of Babesia and Theileria parasites is confirmed to be due to eugenol [27], the main component active against *E. coli* non-growing stationary phase bacteria is unknown and needs to be determined. Allspice is widely used in food processing throughout history and is known to have antibacterial activity against oral pathogens [28], and in this study, its activity against uropathogenic *E. coli* may provide evidence to expand its usage. Bandit "Thieves" is a synergy blend of essential oils containing clove, cinnamon, lemon, rosemary and eucalyptus oils. The anti-persister activity of Bandit "Thieves" may depend on the dominant components including cinnamon bark, clove oil which each has been shown to have activity against UTI89 in this study and their synergistic effect. The attractive bioactivity of this blend Bandit "Thieves" also gives a hint on using oil combinations or oil antibiotic combinations in future studies for developing possible more effective treatments.

Although Health shield, Cinnamon leaf and Clove bud could not clear all *E. coli* persisters at low concentrations (0.5% and 0.25%), they showed obvious activity at a high concentration (1%) with 100% clearance after just one-day essential oil exposure. Meanwhile, they had the same low MIC value (0.125%), which presents strong activity against log phase *E. coli*. Health shield is a synergy blend of essential oils containing cassia, clove, eucalyptus, lemon and rosemary. Compared with Bandit "Thieves", another synergy blend of essential oils containing clove, cinnamon, lemon, rosemary and eucalyptus oils, Health shield showed lower activity against both log phase and stationary phase *E. coli*. This could result from additional active essential oil cinnamon in Bandit “Thieves” that is not in Health shield that contributes to higher activity of Bandit "Thieves”. Cinnamon leaf has been shown to have antimicrobial activity against various foodborne pathogens and has a wide use in fresh-cut produce [29]. Clove bud is a natural product with great potential use in food preservation [30]. Along with Cinnamon bark, Cinnamon leaf and Clove bud showed strong antibacterial activity presumably due to the major component eugenol [25, 29, 30]. Some studies demonstrated that eugenol could denature cellular proteins and change cell membrane permeability when interacting with growing bacteria [30], which could indicate it has similar effect against stationary phase bacteria.

In conclusion, this is the first study of high-throughput essential oil screening against stationary phase uropathogenic *E. coli* in vitro with 140 essential oils and identified several potent essential oils. The top hits are Oregano, Bandit "Thieves", Cinnamon bark, Syzygium aromaticum and Allspice. Meanwhile, we conducted the in vitro drug combination study using essential oil (Oregano) and found some effective combinations to kill *E. coli* persisters. Further studies should be carried out to identify the active components, evaluate safety and pharmacokinetics and efficacy properties, and also their ability to eradicate persistent UTIs in the mouse model.

## 4. Materials and Methods

### 4.1. Bacterial strain

UTI89 is a uropathogenic *E. coli* strain isolated from a bladder infection of a woman [8]. The strain was incubated in LB broth overnight to stationary phase without shaking at 37°C, 5% CO2. The stationary phase *E. coli* culture (~10^9^ CFU/mL) was used directly without dilution for essential oil screen and drug exposure.

### 4.2. Antibiotics and essential oils

Tosufloxacin, levofloxacin, ciprofloxacin, nitrofurantoin, sulfamethoxazole, trimethoprim, meropenem, cefdinir and fosfomycin were purchased from Sigma-Aldrich (St. Louis, MO, USA) and dissolved in dimethyl sulfoxide (DMSO) or H2O to form stock solutions. All antibiotic stocks (except DMSO stocks) were filter-sterilized by 0.2 µm filter and stored at −20°C.

Commercially available essential oils were purchased from Plant Guru (NJ, USA), Plant Therapy (ID, USA) and Natural Acres (MO, USA). Essential oils were dissolved in DMSO at 5% (v/v). The 5% essential oils were further diluted with the stationary phase or log phase culture to achieve desired dilution in the following experiments to evaluate their activity against *E. coli*.

### 4.3. Screening of essential oils for their activity against stationary phase *E. coli* UTI89

To evaluate the effect of essential oils on stationary phase bacteria, these essential oils and drugs were added to the 96-plates containing stationary phase bacteria, leaving the first and last columns in each plate for control. In the primary screen, each essential oil was assayed at three concentrations: 0.5%, 0.25% and 0.125% (v/v). Tosufloxacin, levofloxacin, ciprofloxacin, nitrofurantoin, sulfamethoxazole, trimethoprim, meropenem, cefdinir and fosfomycin were used at 50 µM as control antibiotics. The plates were incubated at 37 °C, 5% CO_2_ without shaking. After one or three days of exposure to essential oils and drugs, the bacterial suspension was transferred to LB plate with a 96-pin replicator to monitor the bacterial survival and regrowth. All tests were run in triplicate.

### 4.4. Antibiotic susceptibility test

The minimum inhibitory concentrations (MICs) were determined using microdilution method according to the CLSI guideline [31]. Essential oils were 2-fold diluted from 1% to 0.0075%. Fosfomycin was 2-fold diluted from 512 µg/mL to 0.25 µg/mL as a control antibiotic. The 96-well plates were sealed and incubated at 37 °C overnight without shaking. All experiments were run in triplicate.

### 4.5. Validation of active essential oils by colony forming unit (CFU) count

The stationary phase bacteria were transferred into 2 mL Eppendorf tubes. Essential oils were added at the concentrations of 1%, 0.5% and 0.25%. Tosufloxacin, levofloxacin, ciprofloxacin, nitrofurantoin, trimethoprim-sulfamethoxazole, meropenem, cefdinir and fosfomycin were added to bacterial suspensions at the final concentration of 20 µM, respectively. At different time points, 100 µL bacterial suspensions were collected by centrifugation, washed and resuspended in PBS. After serial dilutions, 10 µL of each dilution was plated on LB plate for CFU count.

### 4.6. Drug combination assay on stationary phase *E. coli* UTI89

Our previous work with Borrelia burgdorferi indicated that certain drug combinations could eradicate non-growing persister bacteria more effectively than single drugs [32]. Based on our results, Oregano had high activity against stationary phase *E. coli*. Thus, in this study, we used Oregano as the common element to test the activity of various two-drug combinations in killing *E. coli* UTI89 stationary phase cells. We evaluated tosufloxacin, levofloxacin, ciprofloxacin, nitrofurantoin, sulfamethoxazole, trimethoprim, meropenem, cefdinir and fosfomycin at the final concentration of 5 µg/mL in combination with Oregano (0.02%). The designed drug combinations or single drug controls were added directly to stationary phase culture and CFU count was performed after one-day, three-day, five-day and seven-day drug exposure.

## Author Contributions

Conceptualization, Y.Z.; Data curation, S.X.; and W.S.; Formal analysis, S.X.; and P.C.; Funding acquisition, Y.Z.; and S.X.; Resources, Y.Z.; Writing—original draft preparation, S.X.; Writing—review and editing, Y.Z.

## Acknowledgments

We acknowledge Xiao Ma for discussion on essential oils preparation. We think Rong Quan for helping on *E. coli* culture.

